# Using targeted genome integration for virus-free genome-wide mammalian CRISPR screen

**DOI:** 10.1101/2020.05.19.103648

**Authors:** Kai Xiong, Karen Julie la Cour Karottki, Hooman Hefzi, Songyuan Li, Lise Marie Grav, Shangzhong Li, Philipp Spahn, Jae Seong Lee, Gyun Min Lee, Helene Faustrup Kildegaard, Nathan E. Lewis, Lasse Ebdrup Pedersen

## Abstract

Pooled CRISPR screens have been widely applied in mammalian cells to discover genes regulating various phenotypes of interest. In such screens, CRISPR components are generally delivered with a lentivirus. However, lentiviral CRISPR screens are limited by unpredictable genome insertion, the requirement of biosafety level II lab facilities and personnel trained to work with viruses. Here we established a virus-free (VF) genome-wide CRISPR screening platform for Chinese hamster ovary (CHO) cells with 74,617 gRNAs targeting 18,353 genes. Each gRNA expression cassette in the library is precisely integrated into a genomic landing pad thus reducing the clonal variation. Using this VF CRISPR screening platform, 338 genes are identified as essential for CHO cell growth and 76 genes were found to be involved in the unfolded protein response (UPR) induced endoplasmic reticulum (ER) stress. Extensive validation of the candidate genes further demonstrated the robustness of this novel non-viral CRISPR screen method.

## MAIN TEXT

CRISPR screens have been widely applied to decipher mammalian gene function at genome-scale but generally rely on lentiviral delivery methods^1–3^. Using a viral vector to perform CRISPR screens comes with several limitations^4–6^ such as: (1) Lentiviral libraries show unpredictable insertion locations in the genome leading to variable gRNA expression and possible disruption of endogenous genes. (2) Viral integrations result in varying copy numbers per cell. (3) In most countries, working with viruses requires specialized facilities and trained personnel (4) Lentiviral screens are unsuitable for clinical purposes where a viral vector is prohibited e.g. stem cell therapy.

To overcome these shortages, a few studies have reported virus-free (VF) CRISPR screen systems in mammalian cells. These include replacing an integrated dummy gRNA with a pooled library of gRNAs using homologous recombination^7^, or measuring gene mutation by transient CRISPR-mediated knockout (KO) using whole-genome sequencing^8^. However, these strategies suffer from various bottlenecks: (1) Homologous recombination efficiency of the gRNA cassette is very low and it cannot be enriched by resistance gene selection, due to risk of random integration. (2) Unexpected gene disruption may result following transient expression of multiple gRNAs from the gRNA library donor plasmid (Supplemental Fig.2a). (3) The cost is high since genome-wide sequencing is required when it is not possible to guarantee a single gRNA integration per cell.

Here we present a novel VF CRISPR screening platform and demonstrate it in Chinese hamster ovary (CHO) cells, the most common mammalian cell systems in the biotechnology industry for producing pharmaceutical proteins^9^. Using 1558 RNA-Seq samples from different CHO cell lines and culture conditions, we selected 18,353 expressed genes from the ∼24,000 genes in the CHO genome^10^. Then, we designed gRNAs against the coding sequences of these genes (Supplemental Table S1). These gRNAs were used to build a genome-wide virus-free library (VF-lib) that contained 74,617 gRNAs targeting 18,353 expressed genes in CHO-S cells, together with 1000 non-targeting control gRNAs (Fig. 1a, Supplemental Table S1). We further compared the clone variance of this library with a lentiviral library (lenti-lib) previously established by us using the classical lentiviral CRISPR screen method^11^ (Fig. 1b).

**Figure 1.**
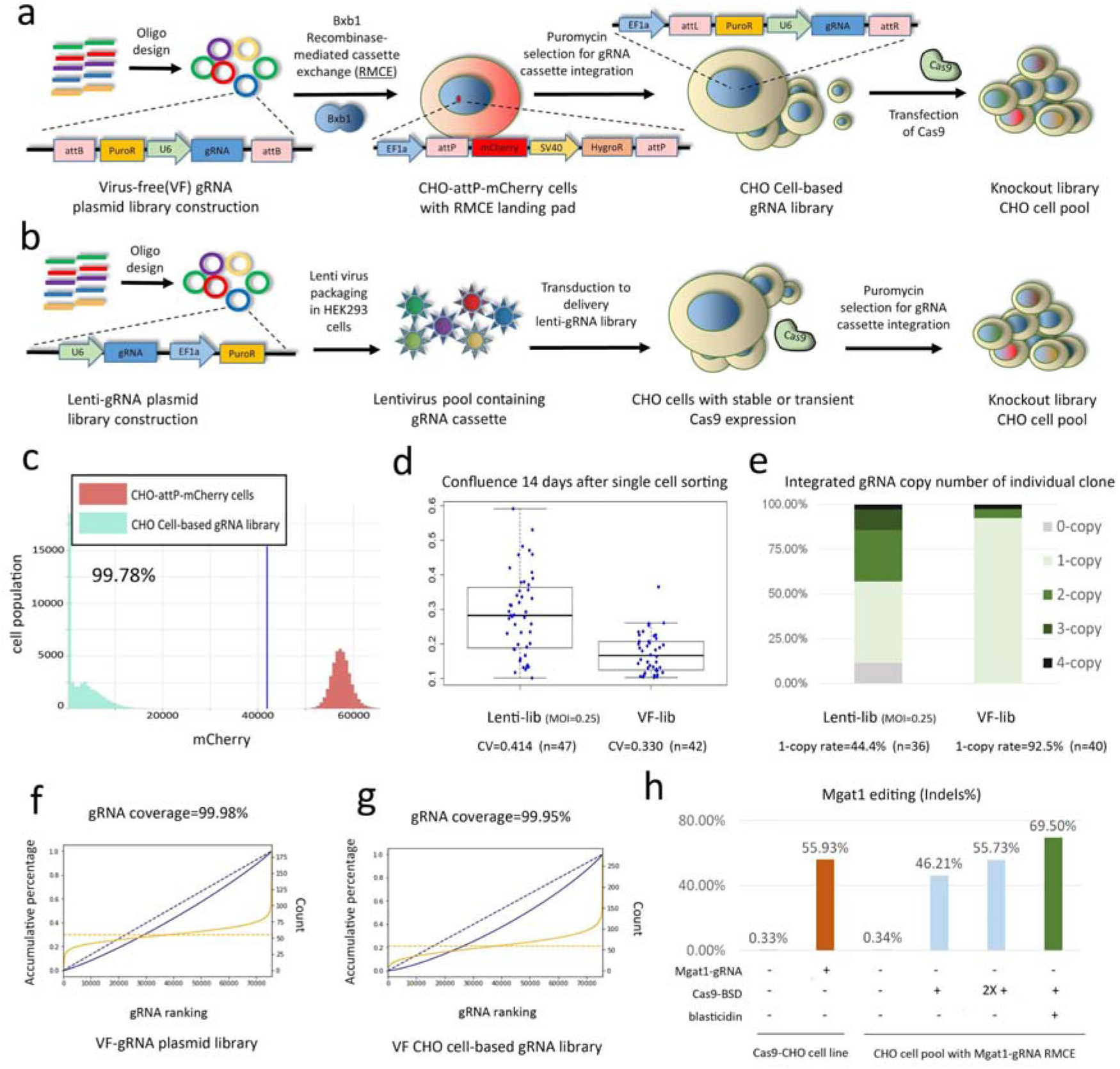
Establishment of a Virus-free (VF) CRISPR screen platform. The illustration of (**a**) VF CRISPR screen and (**b**) lentiviral CRISPR screen in CHO cells. A genome-wide virus-free library (VF-lib) was designed containing 74,617 gRNAs targeting 18,353 expressed genes in CHO-S cells with 1000 non-targeting control gRNAs. The gRNA expression cassette was precisely integrated into the working cell line by Bxb1 recombinase mediated cassette exchange (RMCE). A promoter-less puromycin resistance gene (PuroR) in RMCE donor DNA enables the precise integration of a single gRNA expression cassette. The lentiviral library contained 15,645 gRNAs targeting 2599 genes with 1000 non-targeting control gRNAs (Lenti-lib). (**c**) In the VF-lib, the CHO cell pool containing the gRNA library was enriched after 25 days of puromycin selection. The mCherry cassette was finally replaced by the gRNA expression cassette. (**d**) Before Cas9 was delivered, the cell clone variance in lenti-lib and VF-lib were compared by measuring the colony confluence 14 days after single cell sorting into 96-well plate. CV indicates coefficient of variation. (**e**) The gRNA copy number in lenti-lib and VF-lib was further detected by qPCR. gRNA coverage of VF-lib was calculated in (**f**) the plasmid library after vector construction and in (**g**) the cell-based library 25 days after RMCE of the gRNA library. Cumulative percentage of reads is shown in blue, and the number of reads per gRNA is presented in orange. Solid line = real data, dashed line = ideal model. (**h**) Testing of Cas9 delivery method using *Mgat1* gene as target gene. After establishing a CHO cell pool with RMCE of gRNA targeting at *Mgat1* (Mgat1-gRNA), transient Cas9-BSD vector was transfected. The Cas9-transfected cells were enriched by treatment with blasticidin to kill the cells without transient Cas9 expression. The *Mgat1* editing efficiency was verified by NGS sequencing. Transfection of 1 or 2 rounds (2X +) of Cas9 in the CHO cell pool with RMCE of Mgat1 was set as control. Another method by transfecting Mgat1-gRNA into permanent Cas9 expression CHO (Cas9-CHO) cell line was also compared here.

In the VF screening platform, we used Bxb1 recombinase mediated cassette exchange (RMCE) to precisely integrate a single gRNA expression cassette. We made a CHO-S cell line with a Bxb1 RMCE landing pad that contains an EF1a promoter followed by a recombinase target site (attP), an mCherry expression unit, and an SV40 promoter driving expression of Hygromycin resistance gene (HygroR) followed by a second recombinase target site (mutant attP). The integration site of this landing pad was previously selected as an active site surrounded by highly expressed genes^12–14^. In the donor gRNA library plasmid, the gRNA expression cassette for RMCE is flanked by recombinase target sites (attB and mutant attB). To prevent random integration of the gRNA expression cassette, a promoter-less puromycin resistance gene (PuroR) was designed in the gRNA expression cassette. Therefore, the PuroR gene is only expressed after correct insertion into the landing pad with the EF1a promoter upstream. Lack of mCherry expression thus indicates a successful RMCE of the gRNA cassette. After transfection with the Bxb1 recombinase plasmid and gRNA library plasmids, ∼4% of the cells demonstrated successful RMCE (Supplemental Fig. S1). The RMCE-positive cells were further enriched by puromycin (10µg/mL) selection for 14-20 days (Supplemental Fig. S3). 25 days after gRNA RMCE, the RMCE cell population was fully enriched -- 99.78% cells were negative for mCherry (Fig. 1c). This efficiency is much higher than that achieved previously by another non-viral CRISPR screen approaches that only achieved an efficiency of around 30%^7^.

Undesired gene KOs can be caused by the transient expression of gRNAs immediately after transfection from one or more of the gRNA library donor plasmids prior to genome integration if Cas9 is simultaneously expressed (Supplemental Fig. S2a). This is demonstrated in our preliminary test (Supplemental Fig. S2b-d) and a previous study^7^. We successfully avoided such undesired and hard-to-detect gene editing, by only transfecting with Cas9 25 days after transfecting the gRNA RMCE system to allow sufficient time for dilution/degradation of unintegrated gRNA expression plasmids (Supplemental Fig. S2d).

To investigate the effect of gRNA integration methods on the phenotype of CHO cells, before Cas9 delivery, the variance of cell growth in lenti-lib and VF-lib were analyzed by measuring the colony confluence 14 days after single cell sorting into 96-well plates (Fig. 1d). The cell population in the lenti-lib transduction showed higher variation across cells than the population in VF-lib. We hypothesize that this higher clone variance in lenti-lib is caused by random, and potentially multiple, genomic integrations of gRNA. To investigate the effect of gRNA integration methods on the efficiency of obtaining the desired single copy integration of gRNA cassettes, we used qPCR to compare copy number of genome integrated gRNA cassettes in the VF RMCE cells and cells generated using a traditional lentivirus approach. We found this copy number to be higher and more variable in the cells generated via the lentivirus-based approach compared to the VF RMCE generated cells. The majority (92.5%) of the cells in VF-lib showed single integration of gRNA cassettes, while less than half (44.4%) of the cells in lenti-lib demonstrated such integration events (Fig. 1e). Our previous study also confirmed that 12% (±5%) of cells in lenti-lib contain the integration with more than one category of gRNA which is measured by next-generation sequencing (NGS)^11^. Therefore, the VF integration results in more consistent phenotypes of edited cells in the cell pool before performing the screen.

Due to the efficient and precise RMCE of gRNA cassette in VF-lib, the coverage of gRNA library after integration in the cell-based library was as high (99.95%) as in the plasmid pool (99.98%), and showed an even distribution of gRNAs (skew ratio^1,15^ = 1.85 and 2.84 in the plasmid and cell-based libraries, respectively, Fig.1f, g). To identify an efficient method to deliver Cas9, we tested a vector expressing Cas9 together with blasticidin resistance gene (BSD) in CHO cells followed by treatment of blasticidin to kill the cells without transient Cas9 expression. 3-4 days of blasticidin treatment is sufficient to kill untransfected cells (Supplemental Fig. S4). We also verified this method via NGS by using the *Mgat1* gene as a model. In the blasticidin-selected Cas9-transfected CHO cells with RMCE of gRNA targeting *Mgat1* (Mgat1-gRNA), *Mgat1* gene editing efficiency was ∼70% which is higher than the efficiency via transfection of 1 or 2 rounds of Cas9-BSD without blasticidin selection. Also, the efficiency was higher than transient transfection of the gRNA into a CHO cell line with constitutive Cas9 expression (Cas9-CHO, Fig. 1h). With the Cas9-enriching protocol now validated, we transfected the Cas9-BSD into the cells with the integrated gRNA library, and we subsequently followed by blasticidin selection to establish the final KO CHO cell pool (Supplemental Fig. S4).

9 and 18 days after Cas9 transfection, the genomic DNA from the cell pool was harvested for NGS analysis to measure the gRNA depletion. The percentage of depleted gRNAs increased after Cas9 transfection into the cell-based gRNA library (Fig. 2a). 18 days after Cas9 transfection in cell-based gRNA library, gene set enrichment analysis (GSEA) revealed significantly depleted genes in fundamental cellular processes: genes involved in DNA replication and repair, gene expression, RNA processing and splicing, as well as protein translation^16,17^ (Fig. 2b, Supplemental Table S2). The significant depletion of gRNAs targeting 338 genes was shown in three biologically independent samples compared to non-targeting gRNAs (Fig. 2c, d, Supplemental Fig. S6 and Table S3), indicating that these genes are essential to cell viability or proliferation in our culture conditions. 96% (325/338) of these genes are reported essential either in human cells or in mouse cells^18^ (Fig. 3e, Supplemental Table S4). We further investigated whether the gene essentiality in CHO cells is similar to human cell lines. We analyzed previous pooled CRISPR KO screens in two human cell lines and found that 66% (222/338) and 61% (206/338) of essential genes in CHO cells are also essential in K562 cells^19^ and HeLa cells^20^ (Fig. 2f, g, Supplemental Table S5). Taken together, these results demonstrate that our VF CRISPR screen method can identify essential genes via gRNA depletion analysis.

**Figure 2.**
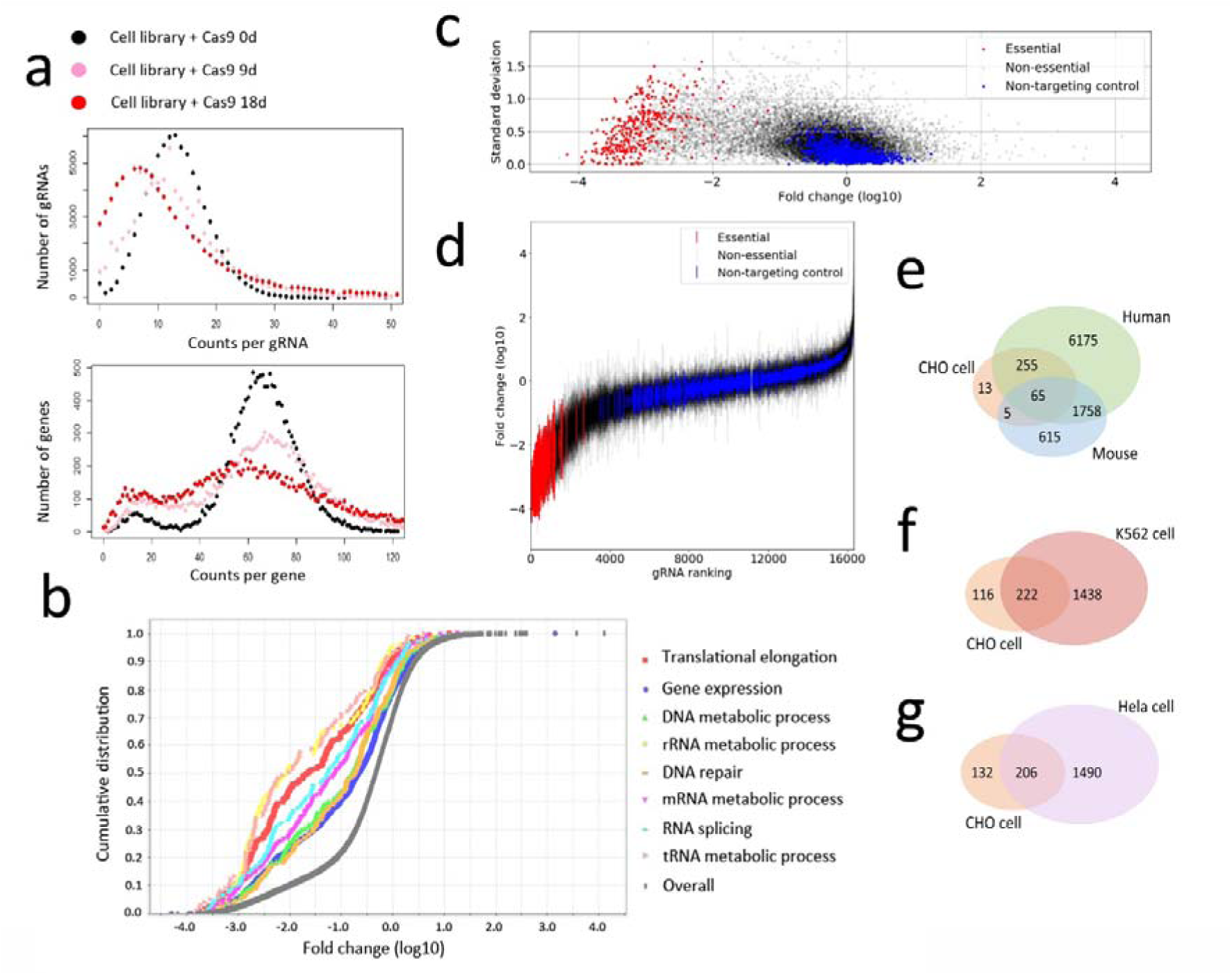
Genome-wide VF CRISPR screen to identify essential genes in CHO cells. (**a**) gRNA reads distribution of targeted genes after Cas9 transfection in the cell-based gRNA library. (**b**) Selected biological processes significantly depleted in CHO cells 18 days after Cas9 transfection in cell-based gRNA library (p<0.05). (**c, d**) Fold change (mean of log10) and the standard deviation of gRNAs among three biological screening replicates. The significantly depleted gRNAs were classified as gRNAs targeting essential genes (highlighted in red). Venn diagram of candidate essential genes in CHO cells and the reported essential genes in (**e**) human and mouse, and in (**f**) K562 and (**g**) Hela cells.

**Figure 3.**
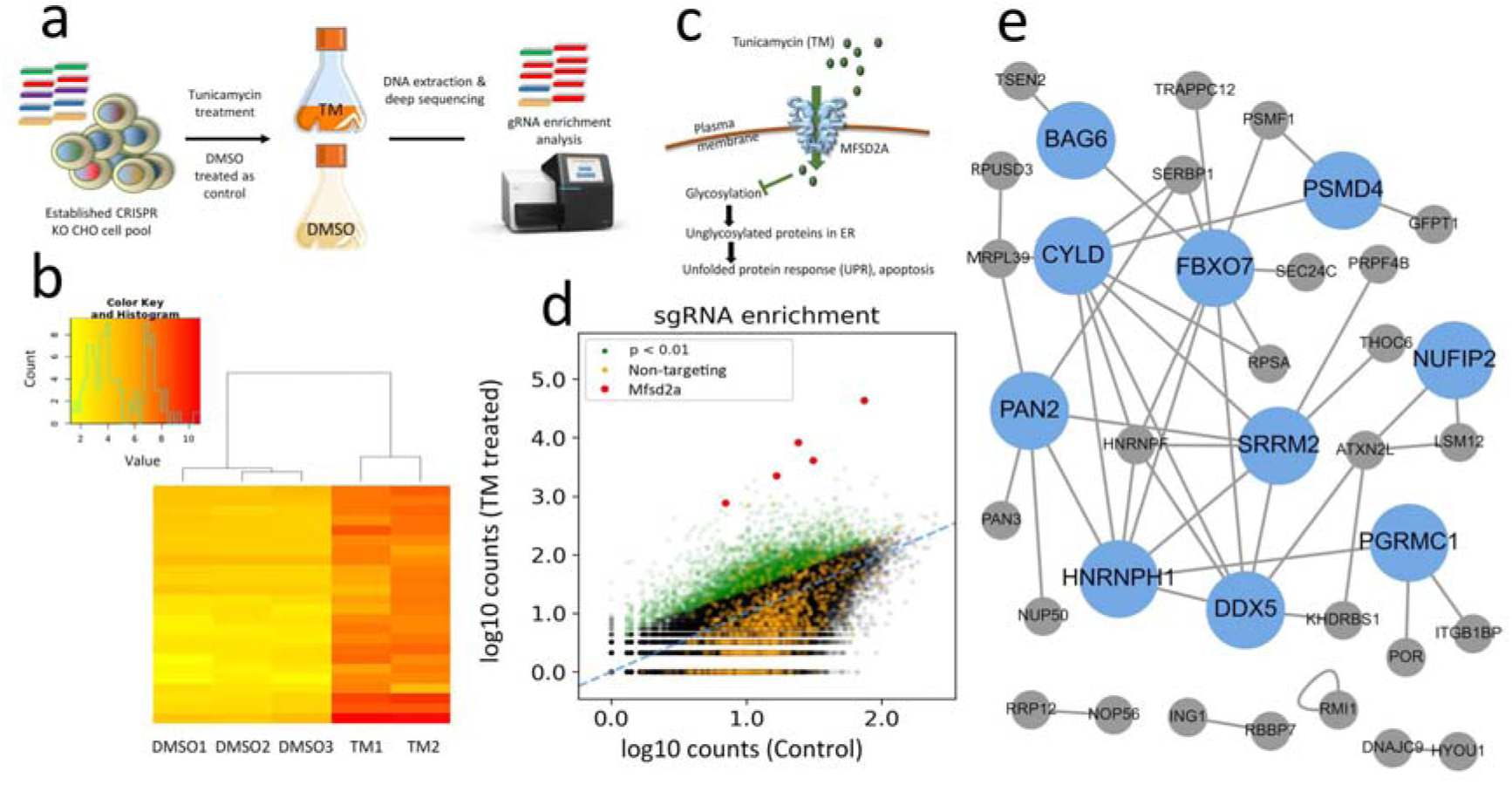
VF CRISPR screen for ER stress resistance genes in CHO cells. (a) Illustration of ER stress resistance cells screen in the CRISPR KO CHO cell pool. ER stress in CHO cells was induced by Tunicamycin (TM) treatment (20ng/mL), while cells treated with DMSO (0.2%) were applied as control. (**b**) Heat map of gRNA enrichment difference in TM treated duplicates and control triplicates. (**c**) Major facilitator domain containing 2A (MFSD2A) transporter as a key mediator in the response to TM. (**d**) gRNAs targeting *MFSD2A* were enriched after TM treatment. (**e**) Network topology-based analysis (NTA) based on the protein-protein-interaction (PPI) of candidate genes whose KO would provide ER stress resistance in CHO cells.

By applying the established VF CRISPR screen platform, we further identified the genes regulating unfolded protein response (UPR) induced endoplasmic reticulum (ER) stress in CHO cells. The CRISPR KO CHO cell pool (60mL, starting density=7×10^5^ cells/mL) was treated with tunicamycin (TM, 20ng/mL) to induce ER stress (Fig.3a). After 4 days of TM treatment, cells were allowed to recover for 5 days by culturing the cells in the medium without selection chemicals (Supplemental Fig. S5). The fold-change of gRNAs in TM-treated and control cells was evaluated by NGS and analysis^21^, and the different fold-change patterns were observed in TM-treated (in duplicates) and control (in triplicates) groups (Fig.3b). *Mfsd2a* (major facilitator domain containing 2A) was chosen as the positive gene control representing the gene KO to obtain ER stress resistance, since *MFSD2A* has been identified as an essential plasma membrane transporter of TM in human cell line^22^ (Fig. 3c). As expected, the gRNAs targeting *Mfsd2a* were significantly enriched after TM treatment (Fig. 3d), indicating the *Mfsd2a* KO cells were resistant to TM-induced ER stress and apoptosis. In total, upon knockout, 76 genes in CHO cells reduced ER stress (combined p-value < 0.01 in Supplemental Table S6). Network topology-based analysis (NTA)^23^ demonstrated that 36 of these candidate genes showed a high degree of protein-protein-interactions (PPI) between candidate genes (Fig. 3e). Further analysis of PPI using the STRING database^24^ demonstrated that these candidates significantly clustered (p = 1.99e-09; Supplemental Fig. S7).

Finally, we selected *Bag6* (BCL2 associated athanogene), *Zfx* (Zinc Finger Protein X-Linked) and *Ddx5* (DEAD-Box Helicase 5) for further validation since all the gRNAs targeting them significantly increased after TM treatment (Fig. 4a-d). In addition to the five original gRNAs in VF-lib, one new gRNA was designed targeting *Bag6, Zfx* and *Ddx5*, respectively (Fig. 4e). 7 days after transfection of the new gRNA for each gene, the different cell pools were treated with TM (20ng/mL) for an additional 4 days. After TM treatment, the Cas9-CHO cells transfected with gRNAs targeting *Bag6* or *Zfx* survived better than the control cells, transfected with non-targeting gRNA (NT-gRNA) or GFP based on their viability and viable cell density (VCD) measurement. For example, the *Bag6* mutant recovered to 88% viability by 4 days after TM treatment (Fig. 4f) and the VCD doubled compared to the control groups (Fig. 4i). Surprisingly, almost full resistance to TM was obtained in CHO cells after disruption of *Zfx*, wherein the cell viability after disruption of *Zfx* remained above 90% (Fig. 4g) and the VCD was almost 5 times more than the control groups (Fig. 4j) 4 days after TM treatment. Interestingly *Ddx5* was identified as an gene could inhibit cell growth (Supplemental Fig. S6, Table S3), however when exposed to TM, *Ddx5* KO appears to have a beneficial effect (Fig. 4h,k), though not as strongly as that observed after disruption of *Bag6* and *Zfx* (Fig. 4f-g, i-j). The results also confirm that disruption of *Ddx5* can reduce the VCD significantly compared to the transfection control in the normal culture medium (Supplemental Fig. S8f), while fewer of such growth-reducing effects was observed after disruption of *Bag6* (Supplemental Fig. S8d). No effect was observed after disruption of *Zfx* (Supplemental Fig. S8e), corresponding to their scoring we measured in essentiality test (Supplemental Table S3). Based on these series of data, the screen score is highly correlated to the gRNA validation, thus the VF CRISPR screen is able to successfully identify candidate genes.

**Figure 4.**
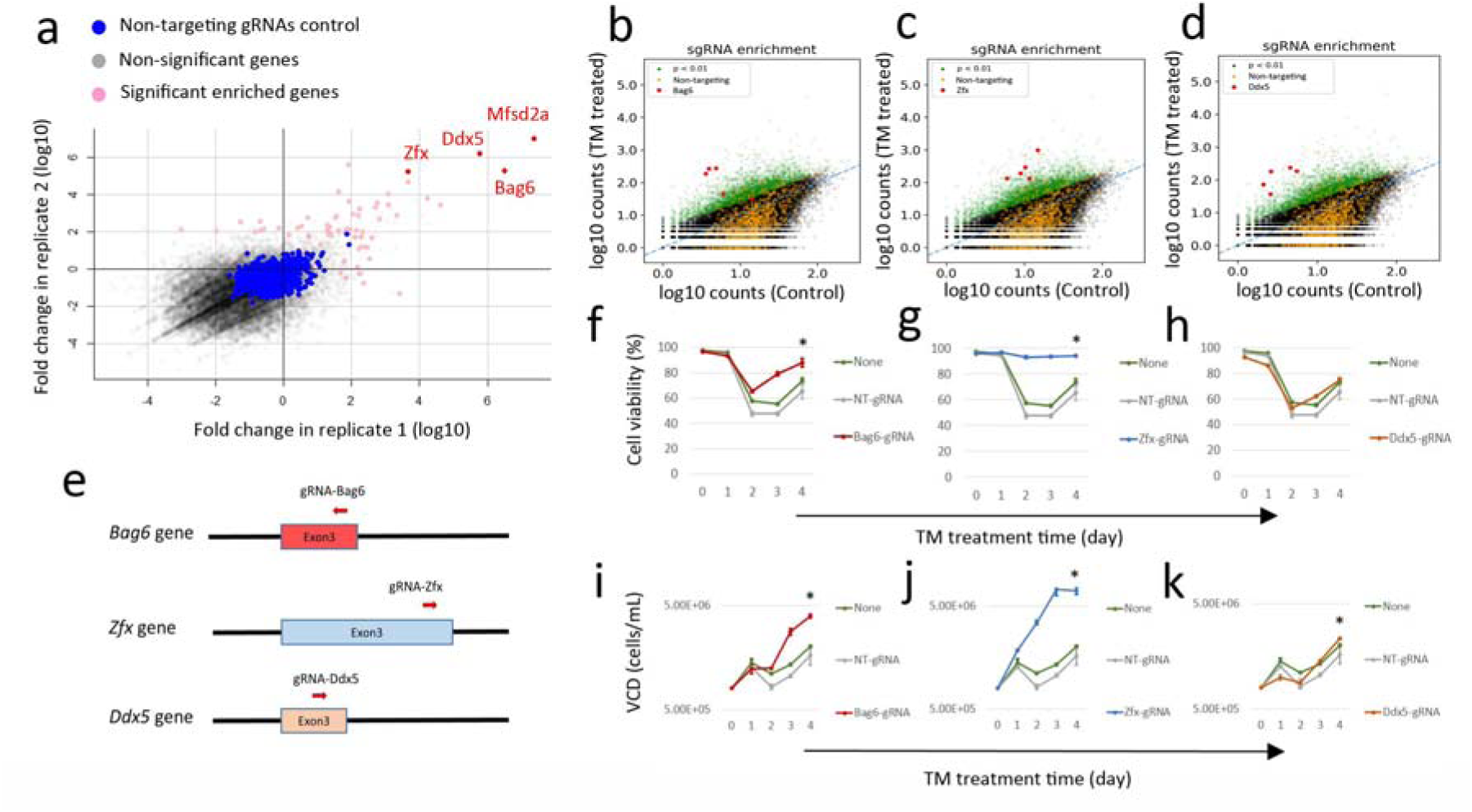
Validation of candidate gene for KO to obtain ER stress resistance. (**a**) Fold change of gRNAs in VF CRISPR screen after TM treatment. Fold change of gRNAs targeting *Bag6, Zfx, Ddx5* and the gRNAs targeting the positive control *Mfsd2a* were highlighted in red. Enrichment of five gRNAs targeting (**b**) *Bag6*, (**c**) *Zfx* and (**d**) *Ddx5*, respectively, in **a**. (**e**) One gRNA for each gene was designed to target Exon3 of *Bag6, Ddx5* and *Zfx* gene in addition to the five gRNAs design in VF-gRNA library, respectively. Effects of gRNAs transfection in Cas9-CHO cells (continuously expressing Cas9) on cell viability (**f**-**g**) and viable cell density (VCD, **i**-**k**) targeting (**f, i**) *Bag6*, (**g, j**) *Zfx* and (**h, k**) *Ddx5* to obtain TM resistance. 7 days after transfection of gRNA, the transfected Cas9-CHO cells were treated with 20ng/mL TM, and the viability and VCD is recorded for 4 days afterward. * indicates statistical significance at day 4 (p<0.05).

Here we presented a novel CRISPR screen approach that provides reduced noise for clearer identification of genes associated with a desired phenotype. Nevertheless, there are some limitations in using this VF CRISPR screen: compared to lentivirus delivered CRISPR screens, our VF strategy requires the introduction of a landing pad in the working cell line, diluting the donor library plasmids, and enriching Cas9-transfected cells. However, the more consistent integration site localization substantially decreases variation across clones in the population^25^ as gRNAs are expressed from a consistent genomic background. Moreover, this method requires no virus facilities which should ease its use in Good Laboratory Practice (GLP) and Current Good Manufacturing Practice (cGMP) facilities. Hence, this new CRISPR screen strategy demonstrates obvious advantages for CRISPR screening of mammalian genes in the genome-wide scale.

## ACKNOWLEDGMENTS

We thank Nachon Charanyanonda Petersen and Saranya Nallapareddy for helping with the FACS and molecular cloning. This work was supported by generous funding from the Novo Nordisk Foundation provided to the Center for Biosustainability at the Technical University of Denmark (NNF10CC1016517 and NNF16OC0021638) and from NIGMS (R35 GM119850, NEL).

## AUTHOR CONTRIBUTIONS

L.P., H.K. and G.L. conceived and supervised the project. K.X., H.K., G.L., N.L. and L.P. designed the experiments. K.X. performed the VF CRISPR screen experiment. K.K, H.H and S.L. (Songyuan) performed the lenti CRISPR screen experiment. H.H. selected the expressed genes for VF genome-wide library. L.G. and J.L. designed the RMCE strategy in CHO cells. S.L. (Shangzhong) designed the genome-wide library. K.X., P.S. and L.P. performed the bioinformatics analysis. K.X. wrote the manuscript with the help of all other authors.

## Notes

### Competing Interest Statement

The authors have declared no competing interest.

