## Supplementary material for "Using targeted genome integration for virus-free genome-wide mammalian CRISPR screen": Supplemental Figures and Legends V2 .docx

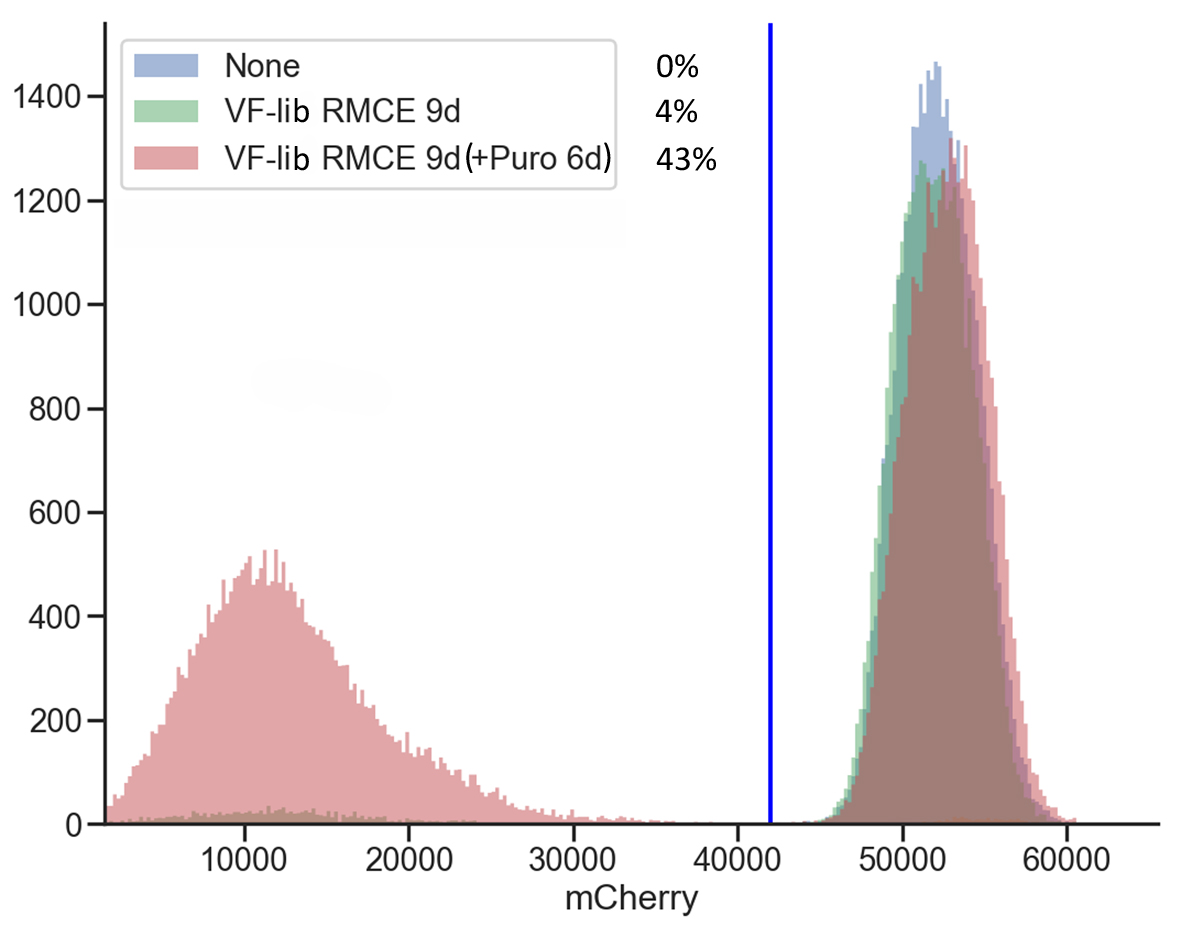

**Figure S1** mCherry density in CHO-attp-mCherry cell lines with or without RMCE of the VF gRNA library. CHO-attp-mCherry cell lines are transfected with the VF-gRNA library and Bxb1 recombinase. After 9 days of transfection, the mCherry density was measured by flow cytometry. The percentage of mCherry negative cells is shown here (mCherry gating = 42000). CHO-attp-mCherry cells without transfection were applied as a control.

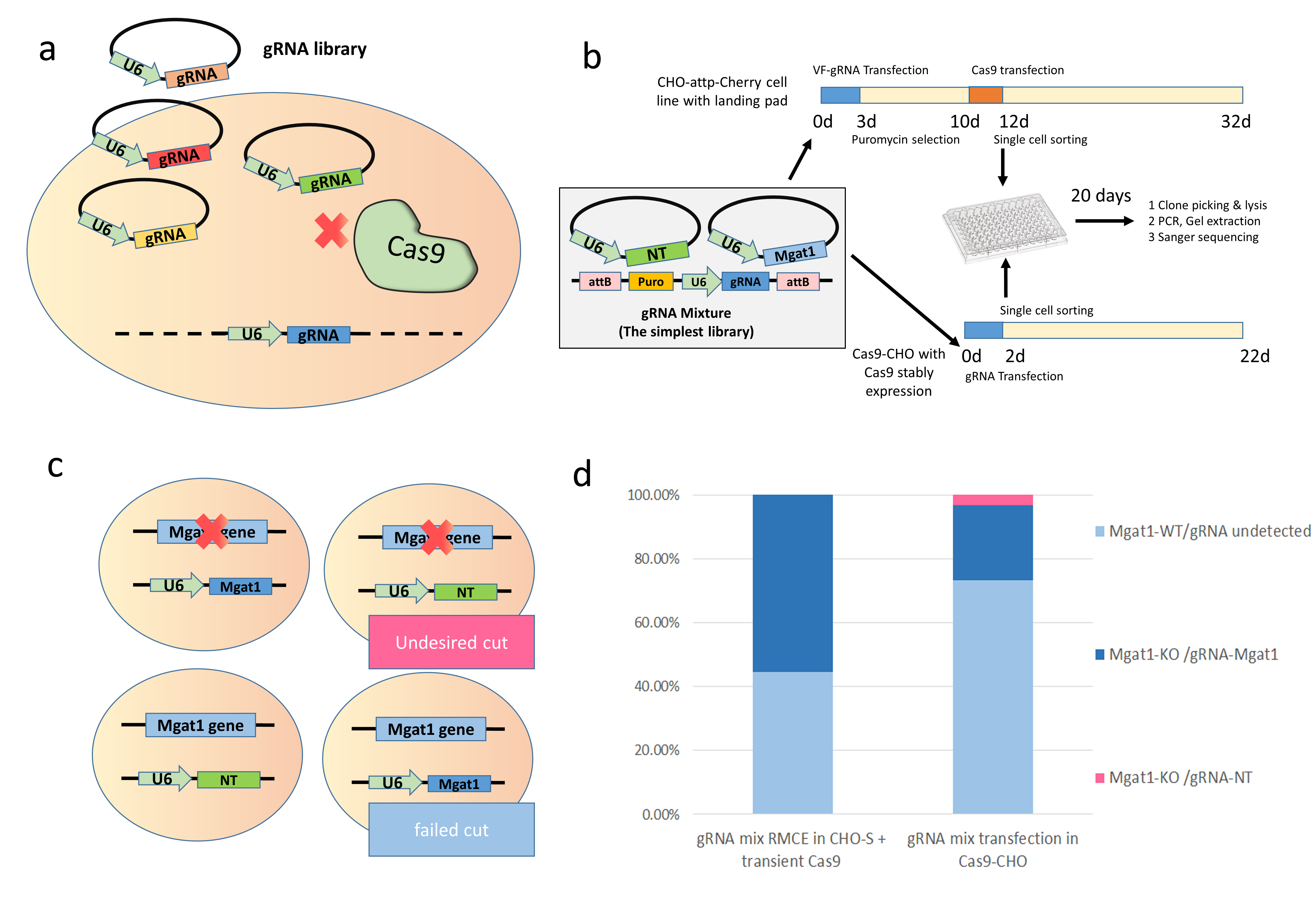

**Figure S2** Undesired gene KO in CRISPR screening while applying a cell line permanently expressing Cas9. (**a**) Illustration of undesired gene editing by the transfected gRNA library plasmid but not by the integrated gRNA cassette. (**b**) Experimental design to compare the effects of undesired gene editing by transfected gRNA plasmid. (**c**)The gene editing and gRNA insertion patterns were compared between the protocol via stably Cas9 expression and the protocol via late transient Cas9 transfection used in this study. (**d**) Percentage of different patterns in two protocols: gRNA mix RMCE in CHO-S + transient Cas9 (n=8), gRNA mix transfection in Cas9-CHO (n=30).

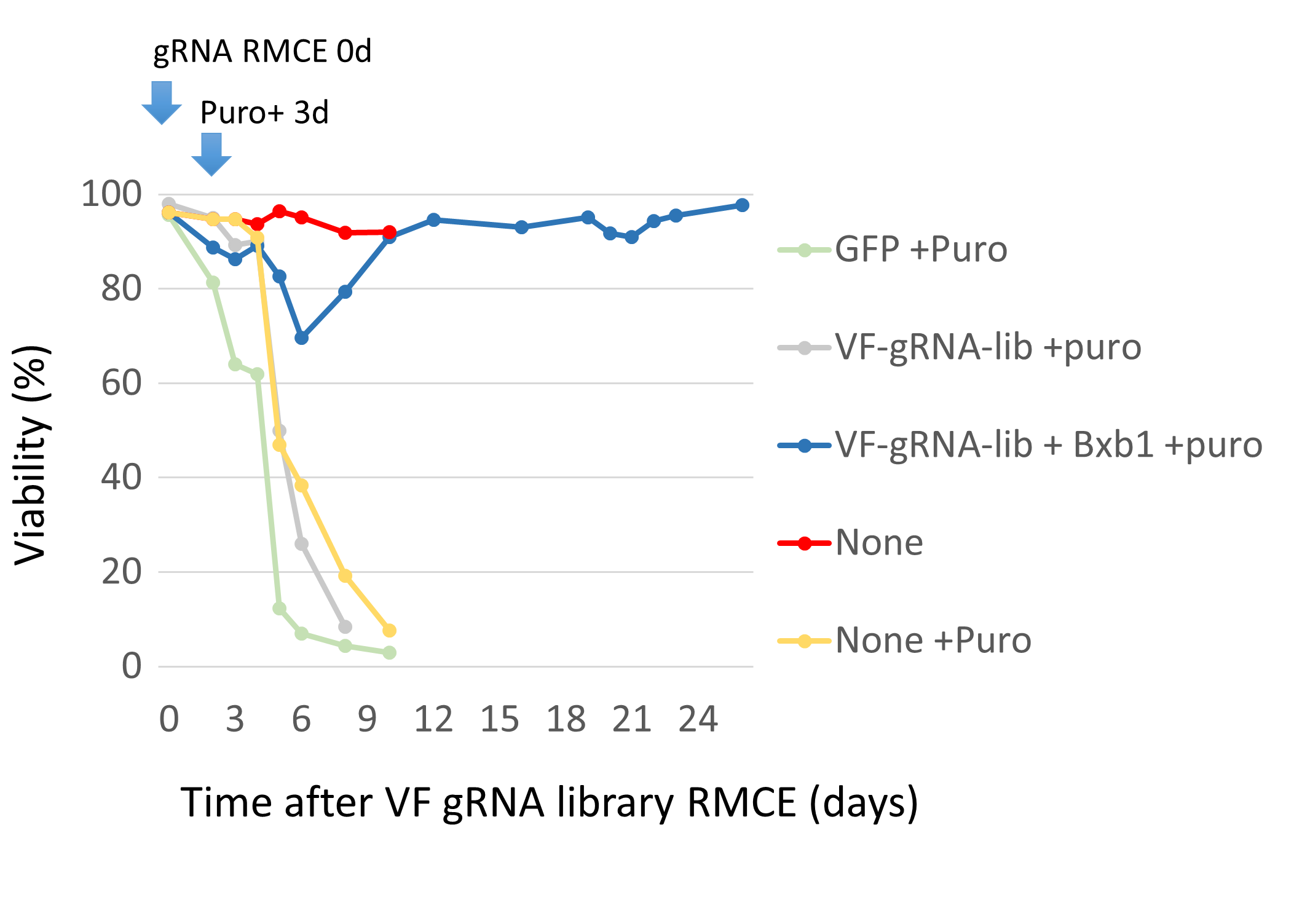

**Figure S3** The cell viability during the process of establishing a VF cell-based gRNA library. CHO-attp-mCherry cells with RMCE landing pads were co-transfected with Bxb1 recombinase and the VF gRNA library. 3 days after transfection, cells were selected with 10µg/mL puromycin for around 20 days. Cells transfected with GFP (GFP + Puro) or without transfection (None + Puro) were set as control for puromycin selection. Cells transfected only with the VF gRNA library but without Bxb1 recombinase (VF-gRNA-lib + puro) confirm the limited random integration of gRNA cassettes. Non-treated cells (None) were used as a control.

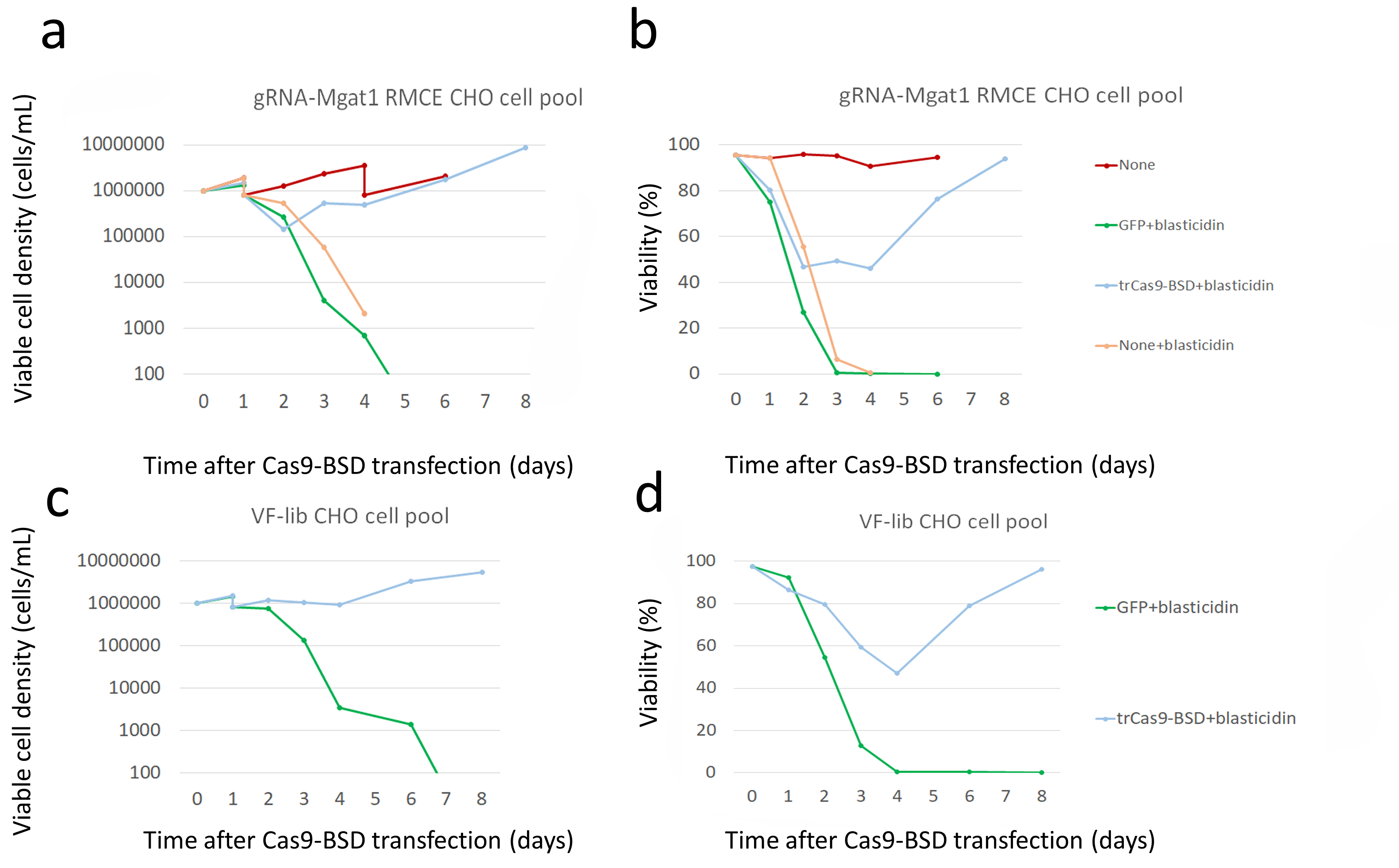

**Figure S4** (**a**) Viable cell density and (**b**) viability of different treated cell pools with RMCE of gRNA-Mgat1 presented in **Figure 1h**. The identical cell pool transfected with GFP was used as a control. (**c**) Viable cell density and (**d**) viability of cell-based gRNA library during the Cas9-transfection enrichment. The cell-based gRNA library transfected with GFP was used as a control. 1 day after transfection of Cas9-BSD, the cells were treated with 10 µg/mL of blasticidin for 1 day and with 5 µg/mL of blasticidin for an additional day. On day 4 after transfection the cells were cultured in medium without blasticidin for recovery. After day 8 the recovered cell pool can be used for furtherexperiments. Cells in the control group would be re-seeded to 1x10^6^ cells/mL if the viable cell density reached 5x10^6^ cells/mL.

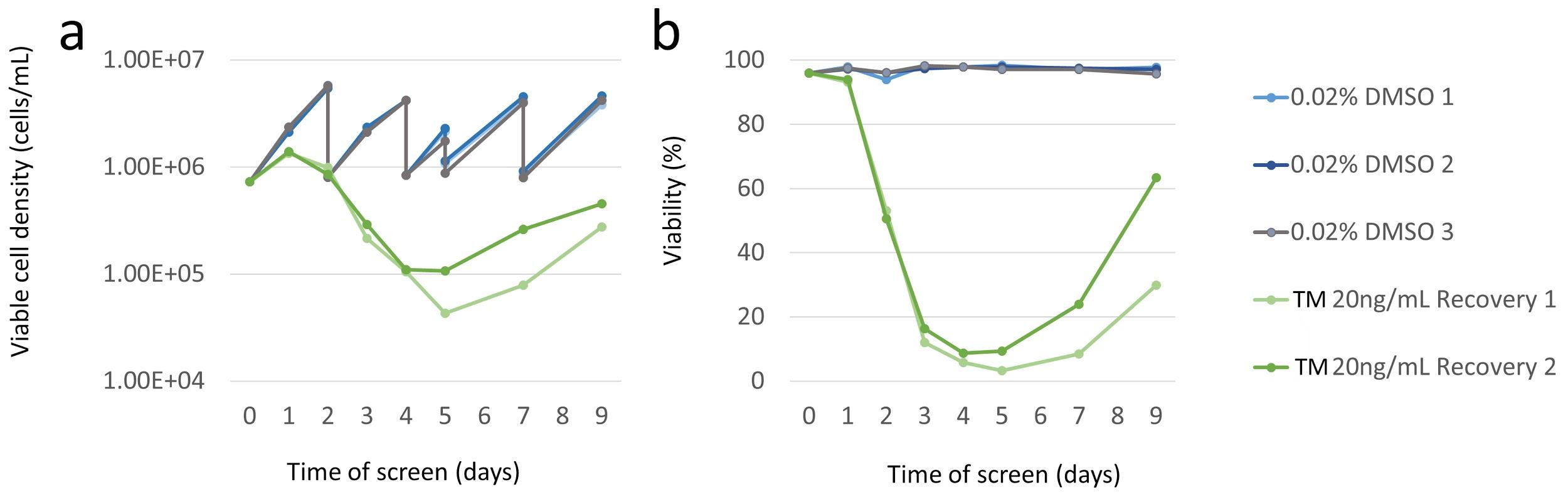

**Figure S5** (**a**) Viable cell density and (**b**) viability of different treated VF CRISPR KO cell pools during ER stress screen. Established VF CRISPR KO cell pools in duplicates were treated with 20ng/mL of TM for 4 days and recovery for an additional 5 days. Cell pools treated with 0.2% DMSO in triplicates were used as a control. Cells in control groups would be re-seeded to 1x10^6^ cells/mL if the viable cell density reached or nearly reach 5x10^6^ cells/mL.

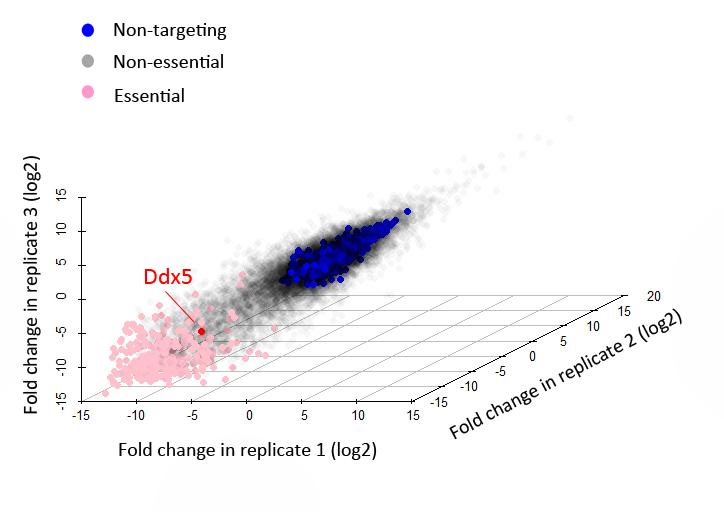

**Figure S6** gRNA fold change among three biological replicates. The genes showing significant fold changes in at least one biological replicate are highlighted as essential genes. The fold change is calculated by averaging the log fold-changes of all gRNAs targeting each gene. The gRNAs targeting these essential genes are demonstrated in pink. The gRNAs targeting *Ddx5* is highlighted in red.

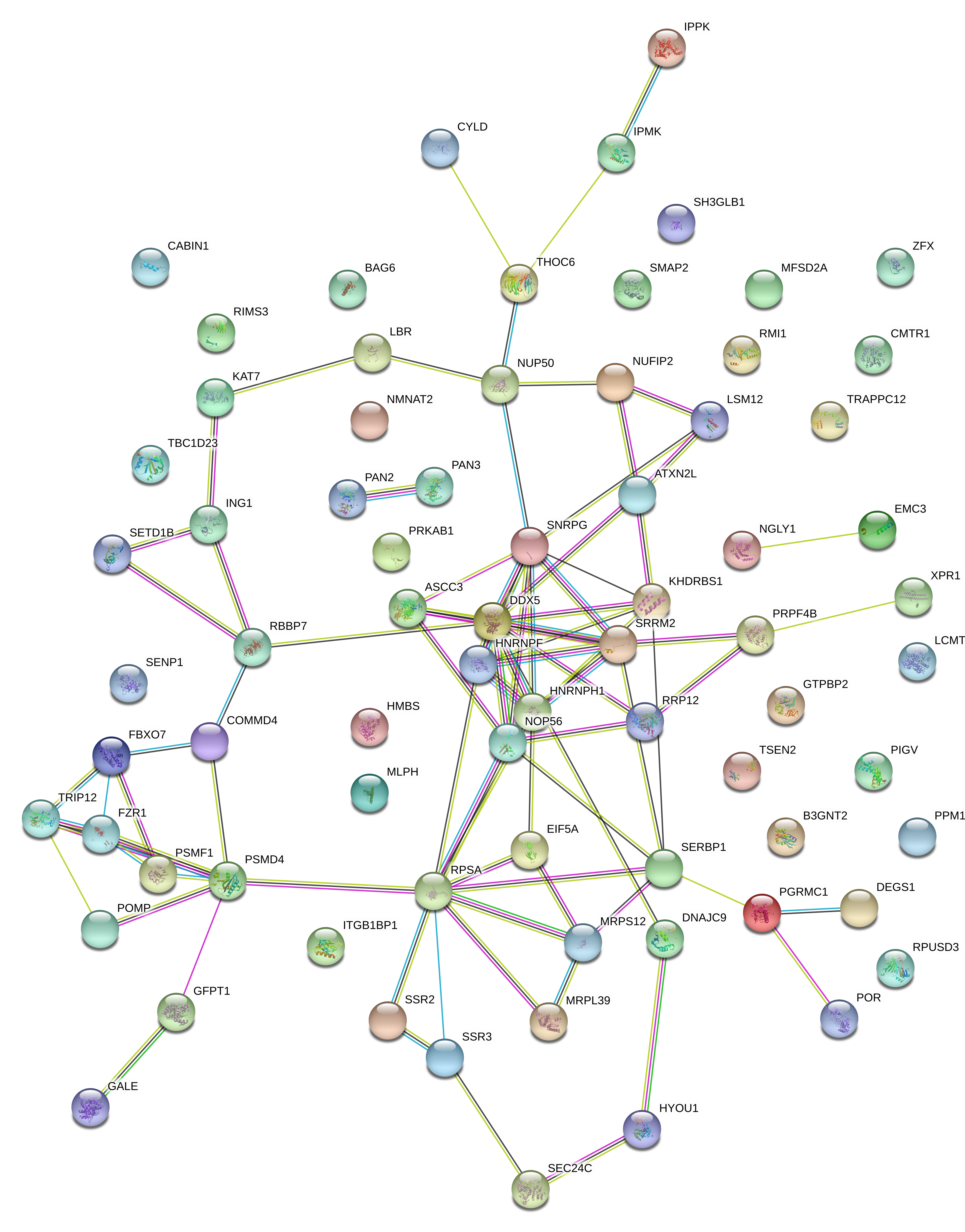

**Figure S7** Protein-protein-interaction (PPI) analysis in STRING database showing the network of candidate genes whose KO would provide ER stress resistance in CHO cells. PPI enrichment p-value = 1.99e-09. p<0.05 means that the input proteins have more interactions among themselves than what would be expected for a random set of proteins of similar size, drawn from the genome. Such an enrichment indicates that the proteins are at least partially biologically connected, as a group.

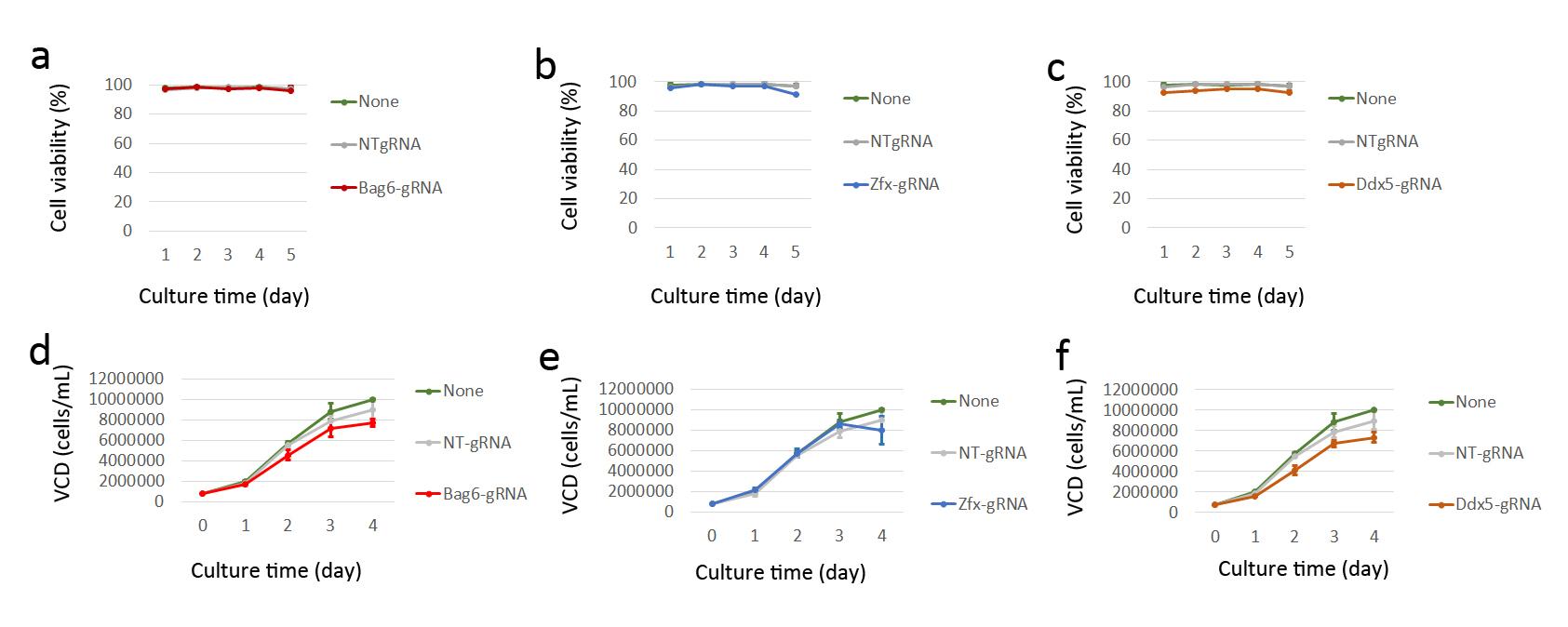

**Figure S8** Effects of gRNAs transfection in Cas9-CHO cells (continuously expressing Cas9) on cell viability (**a**-**c**) and viable cell density (VCD, **d**-**f**) targeting (**a, d**) *Bag6,* (**b, e**) *Zfx* and *Ddx5* (**c, f**). 7 days after transfection of gRNA, the viability and VCD were recorded for 4 days.
